# Comparison of national level spatial and spatio-temporal models of malaria

**DOI:** 10.1101/616847

**Authors:** Kok Ben Toh, Denis Valle

## Abstract

Geospatial statistical models play an important role in malaria control and prevention; they are widely used to produce malaria risk maps, which are essential to guide efficient resource allocation for intervention. Although many models are available for spatial mapping, the most commonly used model in the literature is the Bayesian geostatistical model (BGM), which is based on an underlying Gaussian process. To our knowledge, methods such as splines and decision trees ensemble methods have not been compared relative to their predictive skill for country level malaria prevalence mapping. Moreover, as more countries now have multiple datasets collected throughout the past decade, it is critical to evaluate if the inclusion of past datasets and the use of spatio-temporal models improve the prediction accuracy of present spatial distribution of malaria. Here we compare the prediction accuracy of five models under spatial and spatio-temporal settings in five African countries. The five models are stepwise logistic regression, generalized additive model (GAM), gradient boosted trees (GBM), Bayesian additive regression trees (BART) and the BGM. There is not a single best model to predict malaria prevalence on a national scale. The model performances varied from country to country, and from spatial to spatio-temporal setting. In general, BGM, GAM and BART models performed well, with BGM being the most consistent. The inclusion of past data is not always beneficial: the predictive performance of GAM and GBM increased under spatio-temporal setting, but BGM’s performance decreased in most of the countries. An accurate depiction of malaria risk is important and statistical assumptions that are suitable for a country does not always fit other countries and a wide range of models and settings should be used. It ensures that we find the best modeling approach possible and can provide additional insight to the spatial distribution of malaria risk.

**Author summary:** Malaria is still affecting hundreds of millions of people every year, and killing hundreds of thousands. As the majority of malaria intervention and control policies are developed at the national level, accurate spatial prediction of malaria risk is important. Choosing the best modeling approach for prediction is not straightforward. Here we compare the predictive performance of five models in five countries, with and without dataset from multiple surveys, to provide empirical evidence on whether there is a single best model for national level malaria prediction, and whether inclusion of past dataset may be beneficial in predicting current distribution of malaria risk. We find that models’ performances vary from country to country and there is no single best model. Although Bayesian geostatistical model is widely and commonly used in the literatures, its performance is not necessarily superior to other simpler-to-fit methods such as general additive model (splines) and Bayesian additive regression tree. Importantly, we also show that the incorporation of past data does not always improve spatial predictions of current disease risk. Together, this demonstrate the importance of fitting wide range of models as part of the prediction mapping process, instead of relying on a one-size-fit-all model.

## Introduction

Malaria is still considered endemic in 91 countries today, and continues to devastate people’s health and livelihoods, particularly in Sub-Saharan Africa where 92% of all malaria cases in 2017 were recorded [1]. Malaria is a mosquito-borne disease that is strongly related to socioeconomic and environmental factors. To better understand and more effectively prevent and control the disease, scientists have been leveraging geospatial statistical techniques and increasingly mature remote sensing data products. Currently, there is a large and growing literature on the use of spatial models on malaria at sub-national, national and continental level (e.g. [2–4]). Geospatial statistical models are widely used to identify risk factors, assess efficacy of intervention programs [4], and to produce reliable and comprehensive malaria risk maps [5], which are essential to guide efficient resource allocation for intervention program [6–8].

Bayesian geostatistical models (BGM), which are based on an underlying Gaussian process, are one of the most common type of model in the malaria literature. This technique leverages spatial correlation to improve predictive skill, i.e. it assumes that malaria prevalence among nearby locations is more similar than that of distant locations. This model typically consists of a spatial covariance structure determined by Euclidean distances among sampling points, and a mean function that includes geospatial predictors (e.g. temperatures, rainfall, elevation, vegetation cover, urbanity). This model is fit through Monte Carlo Markov Chain (MCMC) or approximated using stochastic partial differential equation (SPDE) and integrated nested Laplace approximation (INLA) [9, 10]. The latter method is increasingly common due to its speed and computation efficiency, often at a minimal expense of accuracy.

Spatial models of malaria, however, need not be limited to the Bayesian geostatistical model. For example, spline is a common alternative to the Gaussian process regression in spatial models [11]. Although splines have been used to represent nonlinear relationship between geospatial predictors and malaria prevalence [7, 12–14], splines can also be applied to the geographical coordinates to account for spatial correlation. Given an appropriate covariance, spline interpolation is equivalent to kriging [15]; however, their formulation are different, and splines normally produce smoother surface than that of the BGM [16]. Splines can be implemented using Generalized Additive Models (GAM) which is very efficient and fast to fit. Another method that has been gaining popularity in spatial prediction in other fields is the decision trees ensemble method, such as gradient boosted trees, random forests and Bayesian additive regression trees [17–19]. In malaria, this group of methods is uncommon. For example, they have been used to determine the effect of urbanization on malaria prevalence [20] and their predictive performance has been shown to be similar to that of BGM on the sub-continental level [14]. Decision tree ensemble methods allow complicated interaction among geospatial covariates and geographical coordinates, thus potentially accounting for spatial correlation. To our knowledge, the splines and decision tree ensemble methods have not been compared relative to their predictive skill for country level malaria prevalence mapping.

Spatio-temporal models are less commonly employed when modeling malaria prevalence at the national scale, partly because few countries used to have data from multiple years. While this used to be true in the past, more countries now have multiple datasets collected throughout the past decade. For example, the number of African countries with more than one dataset doubled in the last five years according to our review of Demographic and Health Surveys (DHS) datasets. As a result, there is an increasing need to formulate spatio-temporal models at the national level. To this end, modelers can accommodate temporal correlation in various ways, such as the incorporation of an autoregressive process to the existing Gaussian spatial process model or the modelling of spatio-temporal structure with 3D splines in GAM. However, it is unclear if the inclusion of past data and the use of models that account for spatial and temporal correlation necessarily improve prediction of malaria prevalence across the landscape.

In this article, we conduct a model comparison exercise to determine if GAM and decision trees can be good alternatives to the BGM, under both spatial and spatio-temporal setting. We also determine if inclusion of past datasets is beneficial in modeling the current spatial distribution of malaria prevalence. We compared the predictive performance of five modelling approaches in five sub-Saharan African countries (Burkina Faso, Mali, Malawi, Nigeria and Uganda) to find out the best spatial and spatio-temporal models for national level malaria prediction. The five models are stepwise logistic regression, GAM, gradient boosted trees (GBM), BART and BGM, fit using SPDE-INLA. In addition, we compare the performance between the spatial and spatio-temporal models. To help readers use the models discussed here and to reflect on some practical considerations while fitting the models, we have included a short step-by-step tutorial in the Appendix (S1 Appendix) that describes how to implement these models using a simulated dataset.

## Methods

This study is based on Demographic and Health Survey (DHS) and Malaria Indicator Survey (MIS; part of DHS program) conducted between 2009 and 2015 (Table 1). The standard DHS is a household survey on a wide array of population, health and nutrition indicators designed to be representative at national and regional levels (http://www.dhsprogram.com). Typically, a two-stage sampling protocol is followed where a few hundreds “clusters” (e.g. villages or residential areas) are selected with probability proportional to population size and a number of households within each cluster are sampled. Children between six months and five or six years of age (depending on the survey) in a subsample of the selected households are tested for malaria. MIS is similar to DHS but aims primarily at collecting data on malaria indicators. The MIS target population is limited to women of reproductive age (15-49 years old) and children under five years of age. We used the microscopy test outcome as the malaria infection status (0 = negative, 1 = positive). No individual or household identifier was used in this study.

**Table 1.** Description of the dataset used in this study. Prevalence here is unweighted.

| Country | Survey Type | Survey Period | No. of clusters | Sample size | Malaria Prevalence % |
| --- | --- | --- | --- | --- | --- |
| Burkina Faso | MIS [21] | Sep 2014 - Nov 2014 | 248 | 6171 | 49.1 |
|  | DHS [22] | May 2010 - Dec 2010 | 540 | 5741 | 64.9 |
| Mali | MIS [23] | Sep 2015 - Nov 2015 | 177 | 7375 | 35.6 |
|  | DHS [24] | Nov 2012 - Feb 2013 | 408 | 5640 | 50.9 |
| Malawi | MIS [25] | May 2014 - Jun 2014 | 140 | 1982 | 28.6 |
|  | MIS [26] | Mar 2012 - Apr 2012 | 140 | 2121 | 25.1 |
| Nigeria | MIS [27] | Oct 2015 - Nov 2015 | 288 | 5047 | 27.6 |
|  | MIS [28] | Oct 2010 - Dec 2010 | 234 | 5220 | 38.4 |
| Uganda | MIS [29] | Dec 2014 - Jan 2015 | 201 | 4702 | 19.5 |
|  | MIS [30] | Nov 2009 - Dec 2009 | 169 | 3970 | 43.7 |

For each country, the newest dataset is hereby considered the “present” dataset, and the older dataset is the “past” dataset. Under the spatial setting, we used only the present dataset to infer the current (i.e. the newest) spatial distribution of malaria prevalence; under the spatio-temporal setting, we used both present and past datasets to infer the current distribution of malaria prevalence.

All DHS surveys adhere to strict ethical standards. The protocols are reviewed and approved by ICF (the company that implements the DHS program) Institutional Review Board (IRB) and the participating country’s IRB. Informed consent statement is read to the respondents prior to the interview and the biomarker tests, and participation is voluntary. Participation by a child must receive parent or guardian’s consent. Medication and referral to local health facility are provided to all children who are tested positive in the malaria tests. Privacy and confidentiality are maintained during the data collection and processing as outlined in the survey specific reports (See citations in Table 1) and the DHS website (https://www.dhsprogram.com/What-We-Do/Protecting-the-Privacy-of-DHS-Survey-Respondents.cfm). Access to the DHS dataset is only granted for research purpose. Researchers are required to register and provide their research project details before their access to the dataset is granted.

### Geospatial covariates

The geospatial covariates were extracted from various remote sensing and GIS resources that are freely and publicly available (Table 2). They comprise of commonly used socio-economic variables (e.g., urbanity and population density), environmental and climatological variables. All covariates are either long term averages, or based on the nearest available year; with the exception of rainfall and temperature, which are specific to the survey month of the cluster. The value of raster-based covariates were interpolated from the nearest four cells of a given coordinate. Because geographical coordinates are only collected at the center of each cluster, instead of at each household, all individuals within a cluster share the same latitude and longitude coordinates provided by DHS program. Thus, all individuals within a cluster also share the same geospatial covariates. DHS geographical coordinates are randomly displaced to protect the participants’ privacy (two kilometres for urban clusters, five kilometres for rural clusters and one percent of the clusters are displaced up to ten kilometres). Because of the random displacement of geographical coordinates, the extraction of raster data based on a buffer area around each sampled location is recommended [31]. However, we found that the covariates are so highly spatially correlated that point and buffer extraction do not differ significantly.

**Table 2.** Descriptions of the geospatial covariates used in this study

| Covariates | Descriptions | Time | Source |
| --- | --- | --- | --- |
| Population | Number of people per raster cell | 2010 or 2015 | Worldpop [32] |
| Aridity | Aridity index between 0.01 (not arid) to 0.99 (very arid) | 1960 - 1990 | CGIAR-CSI Global-Aridity [33] |
| Built-up Index | Built-up index derived from Landsat | 2014 | GHS built-up grid [34] |
| Enhanced Vegetation Index | Yearly average of the vegetation index between 0 (no vegetation) and 1 (highly vegetated) | The year the survey was conducted | LPDAAC USGS [35] |
| Nighttime Lights | Composite radiance values | 2015 | VIIRS [36] |
| Potential Evapotranspiration | Calculated using WorldClim, mm/yr | 2009 | CGIAR-CSI Global-PET [33] |
| Monthly rainfall | Average monthly rainfall in mm/yr around the survey period | From the two months prior to the month the cluster was surveyed | CHIRPS [37] |
| Elevation | Measured in meter (m) | 1996 | USGS GTOPO30 [38] |
| Monthly Temperature | Average monthly daytime land surface temperature around the survey period | From the two months prior to the month the cluster was surveyed | MOD11B3 [39] |
| Accessibility | The estimated travel time to the nearest high-density urban centres | 2015 | The Malaria Atlas Project [40] |

All covariates, including latitude and longitude, were standardized to have mean of zero and standard deviation of one.

### Models

Five modelling approaches are used in this study. The following sections describe these models and explain how they are formulated for spatial-only (with present dataset only), and spatio-temporal setting (with both past and present data).

In all models under the spatio-temporal setting, the response variable consisted of the microscopy test outcome for each individual. More specifically, let malaria status for individual *i* in cluster *j* at survey year *t* be denoted by *m_ijt_*, where *m_ijt_* = 0 if the child had no malaria (negative microscopy test) and *m_ijt_* = 1 if the child had malaria. All models assume that:

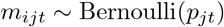

The goal here is to model *p_jt_* (i.e. malaria prevalence of a given location and time) as accurately as possible given the design vector *x_jt_* and, for some models, the spatial-temporal random effects. The design vector contains the predictor variables described in Table 2, latitude, longitude, the interaction between latitude and longitude, and a dummy variable indicating if the observation comes from the present (1) or past (0) dataset. Different assumptions are made regarding the prevalence *p_jt_* in different models.

Under the spatial-only setting, all temporally related components are dropped and all models assume that:

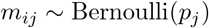

We model the malaria prevalence of given location, *p_j_*, using the design vector *x_j_* and, for some models, spatial random effects. In this setting, the design vector contains the predictor variables described in Table 2, latitude, longitude, and the interaction between latitude and longitude.

### Stepwise logistic regression

The stepwise logistic regression model is the simplest model we adopt and is included as a “baseline” model. It assumes that

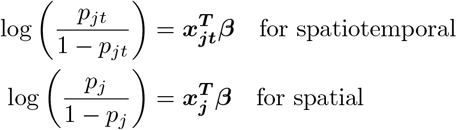

where *β* is the vector of regression coefficients. The regression coefficients are estimated using iteratively weighted least squares. Stepwise regression method was used to select variables based on AIC to avoid the risk of overfitting these data. This is done in R using `stepAIC()` function from `MASS` package.

### General additive model (GAM)

The GAM is similar to logistic regression, with the addition of 2D (involving longitude and latitude) or 3D splines (involving longitude, latitude and year of survey):

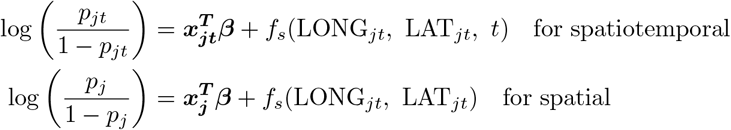

where *f_s_* is a thin plate spline function. This model was fitted using the restricted maximum likelihood (REML) method in the `mgcv` package in R [41]. The package also uses generalized cross validation to determine the optimal amount of smoothness in the spline function.

### Gradient boosted trees (GBM)

Tree-based methods divide the predictor space, i.e. the set of all possible values for the predictors, into *J* parameter regions that are distinctive and non-overlapping. All observations with predictors falling in the parameter region *k* (*R_k_* where *k* = 1, 2,…, *J*) will share the same predicted response. Because a single decision tree is often suboptimal in terms of predictive accuracy, the predictive performance of tree-based method is improved by combining the results of an ensemble of decision trees.

Gradient boosted tree is a popular machine learning technique that is based on an ensemble of decision trees. This method grows the tree sequentially and requires users to predetermine three tuning parameters: number of iterations or number of trees to grow, *B*; the depth or number of splits of each tree, *d*; and a shrinkage parameter or the iterative “learning rate” *λ*. The algorithm starts with a set of initial predicted response 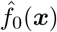. In the *b*-th iteration (where *b* = 1, 2,…, *B*), a new tree *T_b_* with *d* + 1 terminal node is formed to improve the prediction from the previous iteration. This is done by fitting the residuals from the previous iteration to the predictors. The predicted responses of the *b*-th iteration is then updated: 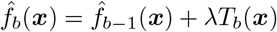. Smaller *λ* or slower learning rate requires more trees to achieve the optimal deviance, and the prediction accuracy is improved from having more trees, although too many trees will eventually have a detrimental impact on the performance. To obtain the predicted response of a given set of predictors 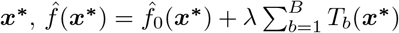.

The fitting process involves gradient descent, thus the method is commonly called “gradient boosting”. Our implementation of the method also uses stochastic subsample of predictors and training samples at each iteration to introduce randomness. Another popular aggregation method is random forest. However, our preliminary study showed that random forest performed much worse than the gradient boosted method across all of our datasets, so we excluded the random forest results from this study.

We implemented the gradient boosting method by using the `gbm` package to fit the model and used the `caret` package [42] to optimize the tuning parameters.

### Bayesian Additive Regression Tree (BART)

The Bayesian Additive Regression Tree (BART) is similar to the Gradient boosted trees in that multiple decision trees are used. BART relies on a probit regression model, in which:

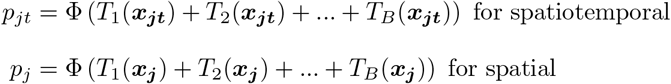

where Φ is the cumulative density function of the standard normal distribution, and *T*_1_,…, *T_B_* are the regression trees. Similar to the gradient boosted trees, model user needs to determine *B*, the number of trees to grow and the structure of the trees is controlled by the prior distribution on tree depth. Finally, strong priors are adopted for the leaf parameters in each tree to ensure the use of multiple decision trees. The model is fit using Monte Carlo Markov Chain (MCMC) and we implement it using the `bartMachine` package [43]. Using cross validation, we determine the optimal number of trees to grow. The model was fitted with 1250 MCMC samples with first 250 discarded as burn-in.

### Bayesian Geostatistical Model (BGM)

Under the spatio-temporal setting, the BGM accounts for spatial and temporal correlations by adding spatio-temporal random effects *α_jt_* to the standard logistic regression framework:

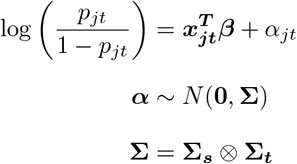

where Σ is the covariance matrix representing the combination of two processes: Σ_*s*_ arises from a 2D Gaussian process that captures the spatial correlation among clusters and Σ_*t*_ arises from a first order autoregressive process (AR1) that describes the temporal correlation between survey years. For spatial correlation, we chose a Matern corelation function of smoothing parameter *ν* = 1, in which the correlation between two clusters *j* and *j*′ at time *t* is:

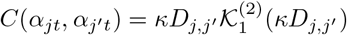

where *κ* is an inverse range parameter, 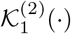 is the modified Bessel function of second kind with order of 1, and *D_j,j′_* is the Euclidean distance between cluster *j* and *j*′. To account for temporal correlation, we assumed an underlying first order autoregressive process, in which the correlation between two different time point *t* and *t*′ of the same cluster *j* is:

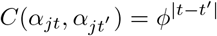

where *ϕ* ∈ (−1, 1) controls the temporal correlation.

Under the spatial-only setting, the BGM formulation above is simplified to:

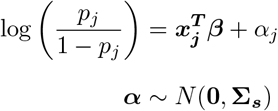

Fitting this model using MCMC is computationally expensive and, as a result, this model is usually approximated by the stochastic partial differential equation (SPDE) approach with a discrete Gaussian Markov Random Field (GMRF; Lindgren et al. 2011). This results in a sparse precision matrix that circumvent repeated and costly matrix inversions. The GMRF structure is determined by SPDE using finite element methods and the resulting model is fit using Integrated Nested Laplace Approach (INLA).

In R, this SPDE-INLA is implemented using the `INLA` package [10, 44]. The target area was first discretized into a 2D mesh of triangles based on sampling location. The mesh was constructed using the `inla.mesh.2d()` function, and the maximum edge length for the inner and outer domain was 0.1 and 1. This results in a mesh of optimum number of fine triangles, i.e. computationally efficient and with unchanged performance as triangle sizes become smaller. The mesh was then used to construct an SPDE model based on the Matern covariance function. Finally, a logistic regression model was fit with all geospatial covariates and a spatial random effect term (i.e. the SPDE object).

### Comparing predictive performance among models

We assessed the predictive performance of the models based on their out-of-sample predictive skill using ten-fold cross validation. Since we are only interested in predicting the “present” prevalence, clusters in present dataset were randomly divided into ten subsets. Data from the first subset of clusters was withheld and the model was fitted using data from the remaining nine subsets, and the past dataset if applicable. Then, we used the estimated parameters to obtain the expected predicted prevalence of the holdout dataset. This procedure was repeated nine times by withholding data from the second to the tenth group of clusters. As a result, each cluster in the present dataset is held out exactly once and has an out-of-sample prediction. We fitted 100 models (five models × two settings×ten folds) for each country.

Using the out-of-sample predictions for each cluster, we calculated the per person log-likelihood (logLik; based on a Bernoulli likelihood) and the population weighted mean absolute error (MAE) based on the mean predicted vs observed prevalence of each clusters. These statistics were calculated using the following expressions:

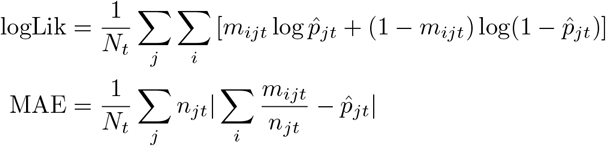

where *N_t_* is the total number of individuals sampled (for the present dataset), *n_jt_* and 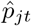 are the number of individuals sampled and the predicted prevalence for cluster *j* at time *t*, respectively.

We compare the performance among models separately for the spatial and spatio-temporal setting. Higher logLik and lower MAE indicate better performance. To determine if a model consistently outperformed another model, we conducted pairwise comparisons to determine the proportion of clusters that have higher logLik in one model compared to another. Because we obtained very similar results when using MAE, we only report the results for logLik for these pairwise comparisons.

Finally, to assess the benefit of using past prevalence data in predicting present prevalence, we calculate the difference in logLik and MAE between spatial and spatio-temporal setting of the same model. If spatio-temporal setting is better, i.e. inclusion of past dataset is beneficial, then we expect that the difference is negative for logLik and positive for MAE.

## Results

### Performance of models for the spatial-only setting

There was no clear “winner” when modelling prevalence using only present dataset (Fig. 1a). The BGM was the best in Burkina Faso and Nigeria (in terms of logLik) and it performed relatively well in all other countries. BART was the best in Mali and in Malawi. It performed well in Uganda but poorly in other countries. GAM was the best model for Uganda, and performed well in Burkina Faso and Uganda. However, its performance in Mali and Malawi was one of the worst among the five models. Interestingly, the stepwise logistic regression performed moderately well in most countries except in Uganda while GBM was the worst in most countries.

**Fig 1.**
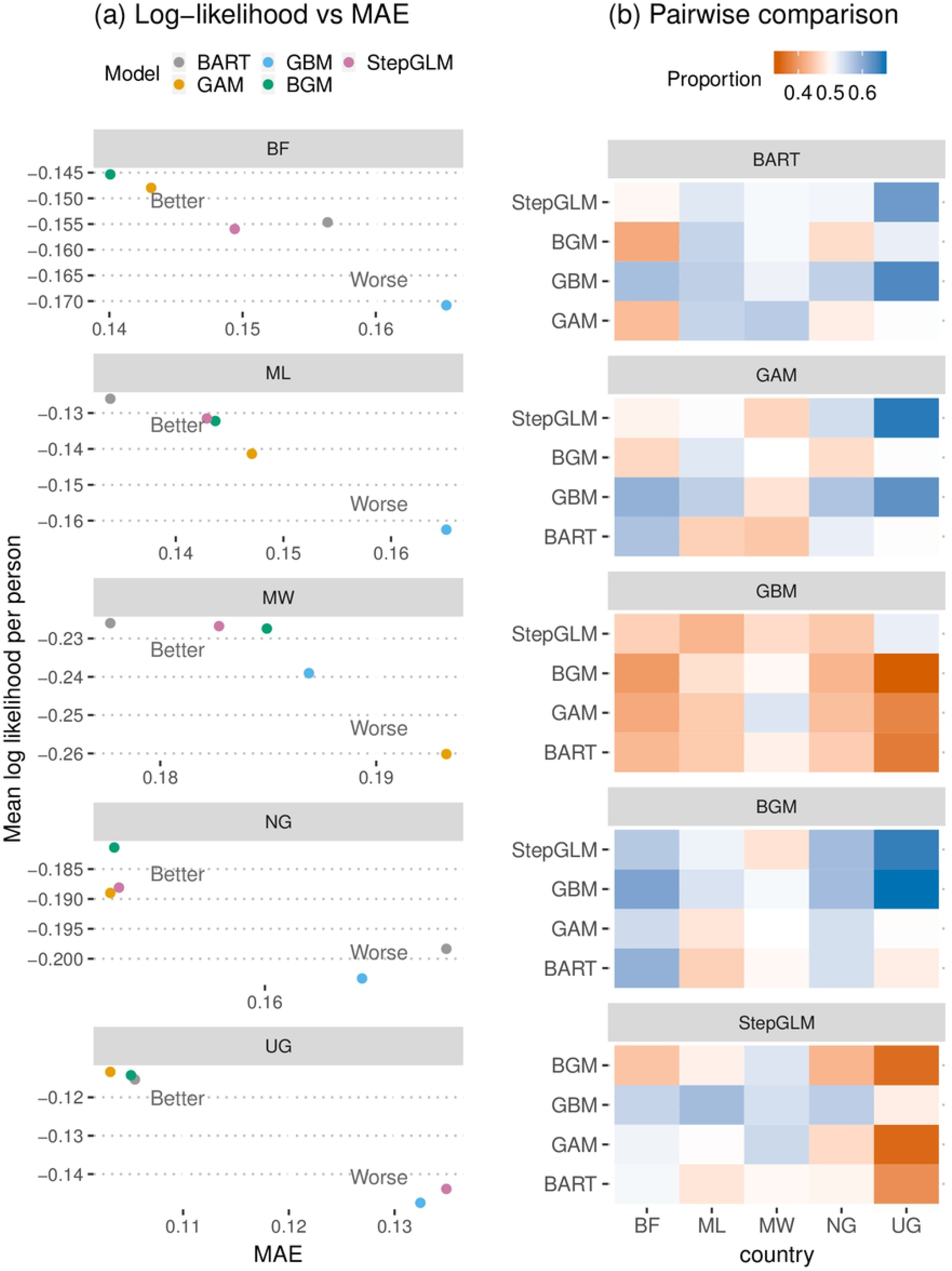
(a) Mean log-likelihood per person (logLik) vs mean absolute error (MAE) for each model under spatial-only setting in each country. Points closer to the top left corner of each panel, i.e. higher logLik and lower MAE, indicate better out-of-sample predictive performance. (b) Matrices of proportion of clusters that a model performed better (based on logLik) than the model on Y-axis under the spatial-only setting. Higher proportion (more blue) means the model associated with the subplot is better and vice versa. StepGLM = Stepwise logistic regression, BF = Burkina Faso, ML = Mali, MW = Malawi, NG = Nigeria, UG = Uganda.

Pairwise comparison among the models yield similar results (Fig. 1b). BGM and BART were the best overall, following by GAM. Interestingly, despite remarkably poor performance in Malawi, BGM was only better than GAM in 50% of the clusters. This suggest that the edge of SPDE-INLA over GAM was not evenly distributed across the country. In general Malawi has the smallest performance difference among models: the maximum proportion of clusters in which one model is better than another was only 0.56 (BART over GAM). The pairwise comparison also confirms that, despite the slightly higher logLik and lower MAE for GAM in Uganda, its performance was tie to BGM and BART.

### Performance of models under spatio-temporal setting

The performances of the models under spatio-temporal setting can be noticeably different from that of spatial setting (Fig. 2a). GAM was the best model in Burkina Faso and Uganda, and second best in Nigeria. However, it remained the worst in Malawi, and its logLik was worst in Mali. BGM’s performance worsened under spatio-temporal setting: it is one of the best models in Nigeria, performed relatively well in Burkina Faso, Malawi and Uganda, but is one of the worst models in Mali. BART’s position was similar to that of spatial setting, but its leading position in Malawi was overtaken by GBM, which is also one of the best models in Mali. However, GBM’s performance in other countries remained poor. The rankings of stepwise logistic regression were poorer in general when compared to spatial-only setting. In Mali and Nigeria, the performance differences among the models reduced remarkably.

**Fig 2.**
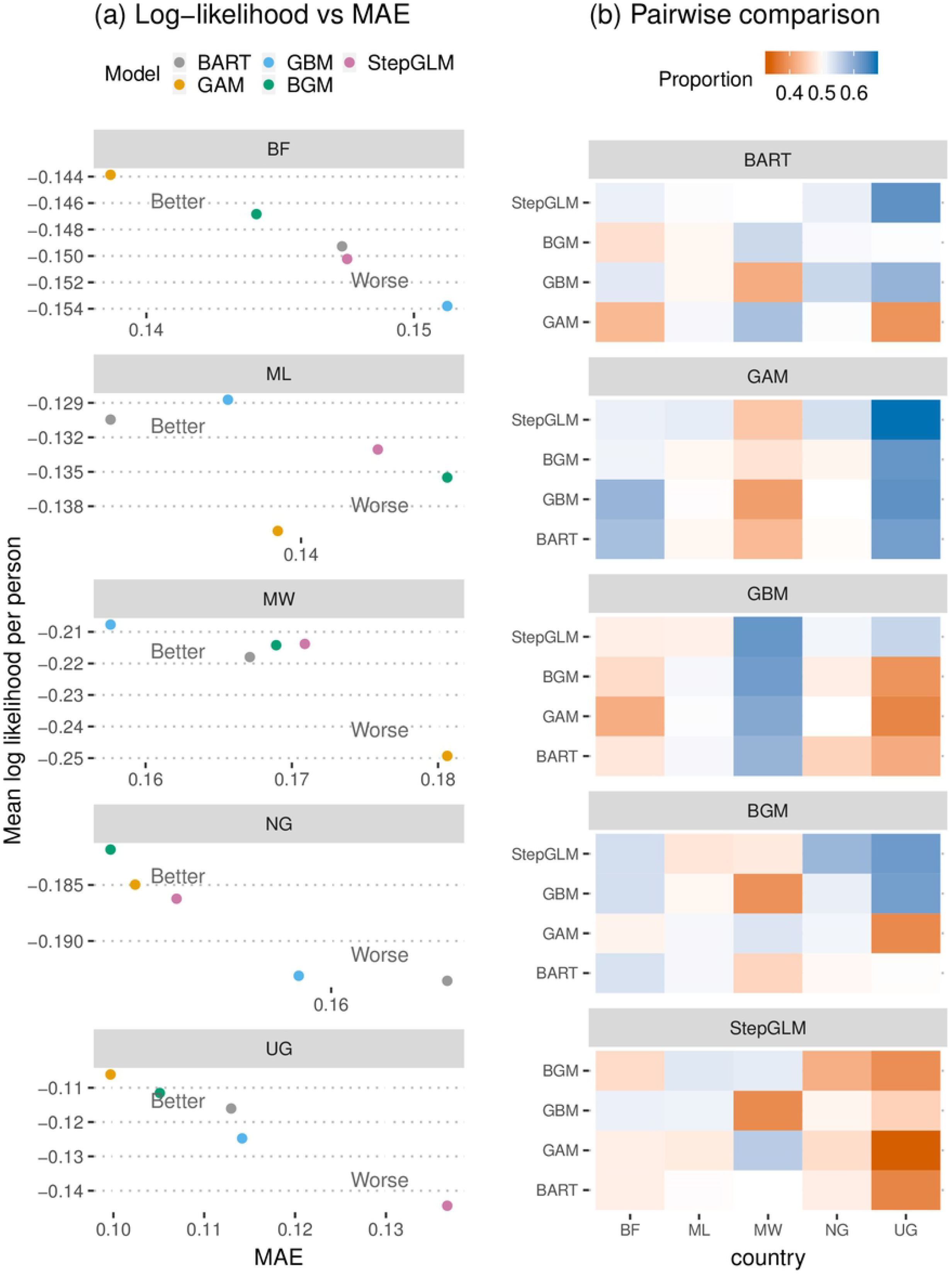
(a) Mean log-likelihood per person (logLik) vs mean absolute error (MAE) for each model under spatio-temporal setting in each country. Points closer to the top left corner of each panel, i.e. higher logLik and lower MAE, indicate better out-of-sample predictive performance. (b) Matrices of proportion of clusters that a model performed better (based on logLik) than the model on Y-axis under the spatio-temporal setting. Higher proportion (more blue) means the model associated with the subplot is better and vice versa. StepGLM = Stepwise logistic regression, BF = Burkina Faso, ML = Mali, MW = Malawi, NG = Nigeria, UG = Uganda.

Pairwise comparison among the models yielded similar results (Fig. 2b). GAM and BGM appears to be the best overall: the former performed better or similar to other models in all countries except in Malawi while the latter noticeably underperformed GBM and BART in Malawi and GAM in Uganda. There were little difference in performance among the models in Mali and Nigeria: the proportion of clusters in which one model is better than another was around 0.52 and 0.53 respectively. On the other hand, GBM appears dominating in Malawi (0.59 to 0.62) while GAM performed much better than other models in Uganda (0.61 to 0.67).

### Performance difference between spatial and spatio-temporal setting

While we expected that having more observations would improve the model performance substantially, our results suggest that the inclusion of past data is not always beneficial when predicting the current spatial distribution of malaria. The use of past data improved the out-of-sample predictive performance of GAM and GBM across all countries (except for GAM’s MAE in Nigeria), but the benefit of including past data is not as evident for Stepwise logistic regression and BART (Fig. 3). Predictive performance of BGM degraded under spatio-temporal setting in the majority of the countries. In Burkina Faso and Malawi, all models (except for the spatio-temporal BGM) performed better with past dataset.

**Fig 3.**
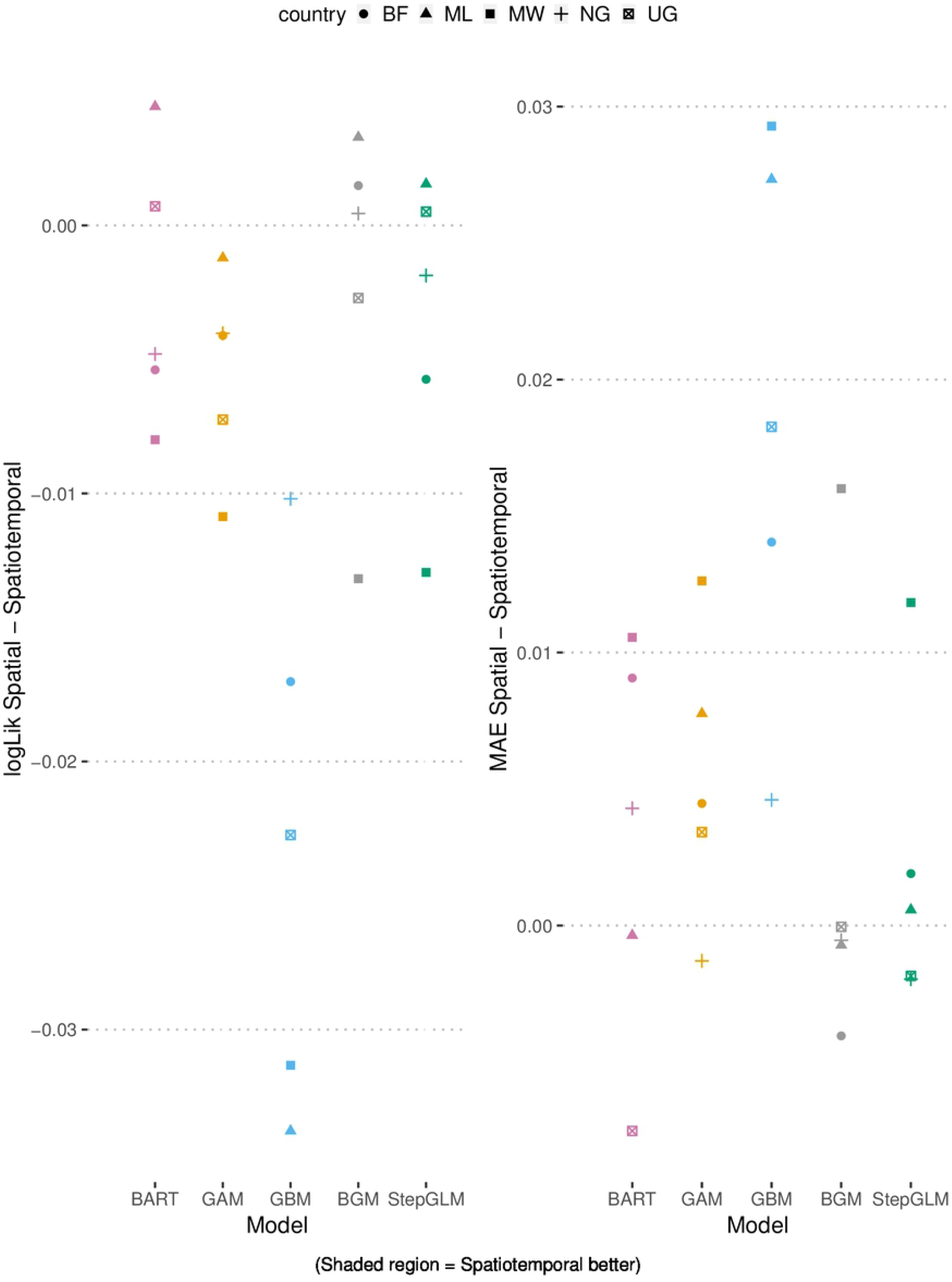
Difference in log-likelihood per individual (logLik) and mean absolute error (MAE) between spatial and spatio-temporal setting of the models. Negative difference in logLik and positive difference in MAE favor spatio-temporal setting. StepGLM = Stepwise logistic regression, BF = Burkina Faso, ML = Mali, MW = Malawi, NG = Nigeria, UG =Uganda.

## Discussion

Based on out-of-sample predictive skill, there is not a single best model to predict malaria prevalence on a national scale and model performance varied from country to country, and from spatial to spatio-temporal setting. However, our results suggest that BGM, GAM and BART are the better performing models, often with little difference in log-likelihood and MAE. Although inclusion of past dataset increases the number of observations, its benefit is mixed: it improves the performance of GAM and GBM, but worsens the BGM in most of the countries.

The Bayesian geostatistical model performs consistently well and is among the top three models across all countries and settings. The model accounts for variation unexplained by the geospatial covariates through the spatio-temporal covariance matrix. However, our separable formulation of the covariance matrix implicitly assumes that the spatial correlation pattern in the past is similar to that of the present. This is a strong assumption that may not be suitable for all countries. For instance, in Burkina Faso where our Bayesian geostatistical model performed remarkably worse under spatio-temporal setting, we observed that the spatial correlation parameter *κ* under spatio-temporal setting is 7.3 times larger than that of spatial setting, i.e. spatio-temporal setting has much lower spatial correlation. For comparison, the ratio of *κ* under spatial-time over spatial setting for other countries was much smaller, ranging from 0.85 in Malawi to 3.07 in Mali. It is possible that the spatial correlation structure of the present dataset in Burkina Faso was very different from that of the past dataset, although the reason for this difference is unclear. Using Gaussian process of non-separable spatio-temporal covariance, might be more appropriate here but this option is not yet available in R-INLA [44].

With the exception of Malawi, GAM’s performed remarkably well, especially under the spatio-temporal setting. Interestingly, despite the ease of fitting and the prediction accuracy, GAM models are rarely used for the spatial mapping of malaria. Our findings somewhat contradict earlier model comparison study which demonstrated that GAM was outperformed by Gaussian process model in all four regions of Africa [14]. The key to this discrepancy is that we have added a spatial (or spatio-temporal) structure to the GAM via the thin-plate spline, and we did not apply the splines to any of the geospatial covariates. Our preliminary results suggest that these steps are critical to the GAM’s success in modelling malaria prevalence, as applying splines on all covariate terms often yielded substantially worse performance, possibly due to overfitting. Unlike our Gaussian process model, GAM uses linear combination of multiple spline terms, instead of a single correlation parameter to describe the spatial structure. Under the spatio-temporal setting, the number of 3D spline terms are about three times the number of 2D spline terms, and the spatial structure of the past is allowed to be markedly different from that of the present. As a result, GAM’s performance is unaffected by inclusion of past dataset with dissimilar spatial structure, and recorded remarkable gain in predictive power due to increased number of observations. It is unclear why GAM’s performance was much worse than other models in Malawi. Since GAM’s spatial or spatio-temporal splines require much more parameters than other models, its performance might have been impacted by Malawi’s low number of clusters in both present and past datasets.

The performance by the decision tree ensemble methods was mixed: GBM was often one of the worst model and BART’s performance was generally moderate. These methods account for spatial-temporal variation by allowing complicated interactions among covariates, which includes longitude, latitude and time. However, this appear to be insufficient in our study and in many cases, GBM was outperformed by a stepwise logistic regression. Our finding is different from that of the model comparison study on the subcontinental level [14], which suggested that GBM’s performance is only marginally behind the Gaussian process model, and is better than GAM and elastic net regularized linear regression. We think that the change in geographical scope and the size of data is the main reason behind the disagreement. GBM’s performance was vastly improved when the dataset is enlarged, i.e. when the past dataset is included in the fitting process. This is particularly evident in Malawi, where number of clusters and number of individuals per cluster in a single dataset is much smaller. Although all models in Malawi improved when we moved from using single dataset to using both past and present datasets, GBM’s improvement was larger and it became the best model under spatio-temporal setting. On the other hand, BART is relatively insensitive to changes in the data size. Its strong performance in Mali and Malawi indicate that it can be a good choice for predicting malaria prevalences when data are limited.

Although the Bayesian geostatistical model, with stationary covariance and linear mean function, is the “standard” model in malaria literature, our study found that it is not always the best performing model. The model fitting can be relatively complicated and it is the most time-consuming method among the models we have used here. Our preliminary study suggests that SPDE-INLA reduces the fitting time drastically at the expense of slight decline in performance compared to MCMC methods. Nevertheless, SPDE-INLA still requires substantial time to fit, especially in country with large land area such as Nigeria because the number of triangle increases considerably. For example, using the same set of parameters to create the triangular mesh, Nigeria had > 25000 triangles while smaller countries such as Malawi had only 6000 triangles. On the other hand, other models require much less time, e.g. fitting time for GAM, GBM and BART under spatio-temporal setting in Burkina Faso was 0.15, 0.02 and 0.36 of the time to fit SPDE-INLA, without including the time for parameter tuning. The fitting procedures for GAM, GBM and BART are also relatively straightforward using the packages in R and it is simple to conduct parameter tuning for GBM and BART through the `caret` package and built-in functions.

As the majority of malaria intervention and control policies are developed at the national level, country-level analyses play an important role in informing policy makers (e.g. [45–47]). An accurate depiction of malaria risk is critical, and model comparison studies like ours provide empirical and useful information to help users understand the different modeling choices and their likely prediction accuracy. We demonstrate that statistical assumptions that are suitable for a country does not always fit other countries. Despite being relatively unexplored for the mapping of malaria prevalence, models such as GAM and BART are promising methods: they are easy to fit and computationally inexpensive, and can often generate spatial predictions of similar quality as those generated by BGM. Although GAM is no new technique, we identified specific formulation (i.e. splining only the spatial coordinates) that is responsible for good predictive performance. We believe that it is important to fit and cross-check a range of models and setting. To this end, we have provided a short tutorial on fitting the models under different settings in R (see S1 Appendix). Adding these steps will not be much more complicated and time-consuming than fitting only Bayesian geostatistical model, however, it ensures that best possible modeling approach is chosen and can provide additional insight to the spatial distribution of malaria risk.

## Supporting information

**S1 Appendix. A short tutorial on fitting the models in R** R code examples and detailed explanations on fitting the five models under the spatial and spatio-temporal settings.

## Acknowledgments

This study would not have been possible without the access to the Demographic Health Surveys (DHS) and other openly available geospatial datasets.

